# Environment and sex control lifespan and telomere length in wild-derived African killifish

**DOI:** 10.1101/2020.09.03.280792

**Authors:** Martin Reichard, Kety Giannetti, Tania Ferreira, Milan Vrtílek, Matej Polačik, Radim Blažek, Miguel Godinho Ferreira

## Abstract

Telomere length is correlated positively with longevity at the individual level, but negatively when compared across species. Here, we tested the association between lifespan and telomere length in African annual killifish. We analyzed telomere length in 18 *Nothobranchius* strains derived from diverse habitats and measured the laboratory lifespan of 14 strains of *N. furzeri* and *N. kadleci*. We found that males had shorter telomeres than females. The longest telomeres were recorded in strains derived from dry region where male lifespans were shortest. At the individual level, we detected a weak negative association between rapid juvenile growth and shorter telomeres in early adulthood. Overall, average telomere length was a good descriptor of telomere length distribution. However, within-individual telomere length spread was not related to any pattern. This substantial variation in telomere length between strains from different environments provides killifish as powerful tool to understand the evolutionarily adaptive value of telomere length.

## INTRODUCTION

Telomeres comprise the ends of all eukaryotic chromosomes and are composed of repetitive DNA sequences and specialized proteins (de Lange 2005). Telomeres protect chromosome ends from illegitimate DNA repair and, through the action of telomerase, from DNA replication-related chromosome shortening (Olovnikov 1973, Greider and Blackburn 1989, Forsyth, Wright et al. 2002). Telomere attrition results in the loss of its protective function and cell proliferation arrest. Critically short telomeres trigger DNA damage responses and genomic instability, a hallmark of aging (López-Otín, Blasco et al. 2013). Apart from the “end-replication problem”, several other factors have been linked to telomere attrition. These include exposure to oxidative damage (Opresko, Fan et al. 2005, Glade and Meguid 2015, Ahmed and Lingner 2018), inflammation (Zhang, Rane et al. 2016), and environmental stressors (Epel, Blackburn et al. 2004, Chatelain, Drobniak et al. 2020). Cells with short telomeres may undergo apoptosis and, if compensatory cell proliferation is no longer available, short telomeres result in cell senescence and inflammation, ultimately disrupting tissue homeostasis (Roos and Kaina 2006, Armanios, Alder et al. 2009, Carneiro, Henriques et al. 2016, El Mai, Marzullo et al. 2020). Impaired tissue homeostasis is at the core of ageing-associated degeneration.

Although not completely understood, telomere shortening in animals, particularly in humans, is a strong promoter of aging-associated degenerative phenotypes (Armanios, Alder et al. 2009, Moskalev, Shaposhnikov et al. 2013). Short telomeres are closely associated with age pathologies such as diabetes mellitus, cardiovascular disease, atherosclerosis, dementia and various cancers (Kota, Bharath et al. 2015, Sethi, Bhat et al. 2016, Aviv, Anderson et al. 2017, Martinez and Blasco 2018). Loss-of-function mutations in genes related to telomere maintenance are responsible for several human syndromes, such as dyskeratosis congenita, aplastic anemia and idiopathic pulmonary fibrosis (Vulliamy, Marrone et al. 2004, Yamaguchi, Calado et al. 2005, Tsakiri, Cronkhite et al. 2007). Similarly, studies using animal models have revealed that mutations in telomerase give rise to age-associated deficiencies and reduction of lifespan (Rudolph, Chang et al. 1999, Henriques, Carneiro et al. 2013, Harel and Brunet 2015, Lex, Gil et al. 2020).

Previous studies have highlighted an association between telomere length and lifespan (Fick, Fick et al. 2012, Heidinger, Blount et al. 2012, Barrett, Burke et al. 2013, Ibanez-Alamo, Pineda-Pampliega et al. 2018). Correlative analyses suggest that, within the same species, individuals with shorter telomeres are prone to disease and die earlier (Salomons, Mulder et al. 2009, Fairlie, Holland et al. 2016, Wilbourn, Moatt et al. 2018), but it remains unclear whether telomere length is an indicator of individual condition (Glei, Goldman et al. 2016). While the protective functions of telomeres are conserved across eukaryotes, telomere shortening with age is not observed in all species (Monaghan and Ozanne 2018, Burraco, Comas et al. 2020). Unlike humans, some animals do not repress telomerase in somatic cells and maintain long telomeres throughout their lives (Sherr and DePinho 2000, Forsyth, Wright et al. 2002). Across species, long telomeres do not correlate with longer lifespans. In fact, longer-lived mammals possess shorter telomeres than short-lived mammals, such as mice and rats (Gomes, Ryder et al. 2011). Instead, the rate of telomere attrition, rather than telomere length per se, appears to correlate with species-specific lifespans (Haussmann, Winkler et al. 2003, Epel, Merkin et al. 2008, Boonekamp, Mulder et al. 2014, Dantzer and Fletcher 2015, Whittemore, Vera et al. 2019).

In humans, women typically have longer telomeres, experience lower rates of telomere attrition and live longer (Gardner, Bann et al. 2014, Lapham, Kvale et al. 2015, Aubert, Baerlocher et al. 2012). While not universal across species, sex differences in telomere length have been reported in most vertebrate taxa studied (reviewed in Barret and Richardson 2011). However, many of these studies report relatively weak effects that fail to be reproduced. It is also not consistent which sex possesses longer telomeres and it is even less clear whether sex-specific telomere length correlates with lifespan or individual condition (Barrett and Richardson 2011, Wilbourn, Moatt et al. 2018, Bauch, Gatt et al. 2020). Hormonal profiles and reproductive stress have been suggested as the causes of differential telomere shortening, but the mechanisms underlying sex differences in telomere length remain largely unexplored (Hartmann, Reichwald et al. 2009, Barrett and Richardson 2011, Graf, Hartmann et al. 2013, Bauch, Gatt et al. 2020).

We performed a comparative experimental study to examine the association between telomere length and lifespan in African annual killifishes from the genus *Nothobranchius*. This group of small and short-lived fishes, and the Turquoise killifish (*Nothobranchius furzeri*) in particular, has recently been established as a model for aging studies. These fish combine the advantages of available resources (reviewed in Cellerino, Valenzano et al., 2016; Hu and Brunet 2018), cheap and convenient husbandry (Polačik, Blažek et al. 2016) and, importantly, links between laboratory studies and natural populations (Reichard and Polačik 2019). We used a set of strains derived from natural populations, which have evolved along a cline of aridity in their natural habitat (Tozzini, Dorn et al. 2013, Blažek, Polačik et al. 2017), to test how variation in lifespan among strains relates to telomere length. As an extension of the association of longer telomeres with shorter-lived species (Gomes, Ryder et al. 2011), we predicted that telomere length would be negatively associated with lifespan across strains.

Within populations, male killifish are shorter lived than females (Reichard, Polačik et al. 2014, Reichard, Blažek et al. 2020), leading to a prediction that females may possess longer telomeres than males. Telomerase expression can be detected in *Nothobranchius* throughout their lives (Hartmann, Reichwald et al. 2009), but telomeres do shorten with age. Hartmann, Reichwald et al. (2009) reported that in a longer-lived strain (MZM 0304), old adults (26 weeks old) had 8% and 27% shorter telomeres in muscle and skin tissues, respectively, compared to young adults (5 weeks old). In contrast, telomeres do not appear to shorten between the ages of 5 and 13 weeks in the short-lived GRZ strain (Hartmann, Reichwald et al. 2009).

We took advantage of the diversity of killifish strains derived from different wild populations and data on their natural habitats. We measured the lifespan of nine *N. furzeri* strains and five strains of the closely related *N. kadleci* (Bartáková, Reichard et al. 2015) in laboratory conditions. We estimated telomere length in 12 of these strains at the time of sexual maturity (4 weeks) and linked them to juvenile growth. Using Telomere Restriction Fragment (TRF) by Southern blot analysis, we tested the statistical distribution of telomere length within individual samples to understand how variability in telomere length, rather than its mean, is associated with juvenile growth and lifespan estimates. Finally, we report telomere lengths for another seven strains of five other *Nothobranchius* species with various lifespans.

## RESULTS

### Lifespan: males from dry-region strains have shorter lifespans

We used strictly standardized conditions to estimate the lifespan of 14 strains from the *N. furzeri*-*N. kadleci* species complex. Across strains, median lifespan varied almost twofold, from 151 days (inbred GRZ strain) to 276 days (Table 1). Overall, males were shorter lived than females (Mixed effects Cox model controlling for population identity nested within species identity: z = 5.39, p <0.001). This pattern was especially consistent in *N. furzeri*. Sex-specific lifespan estimates for each strain are detailed in Table 1.

**Table 1.**
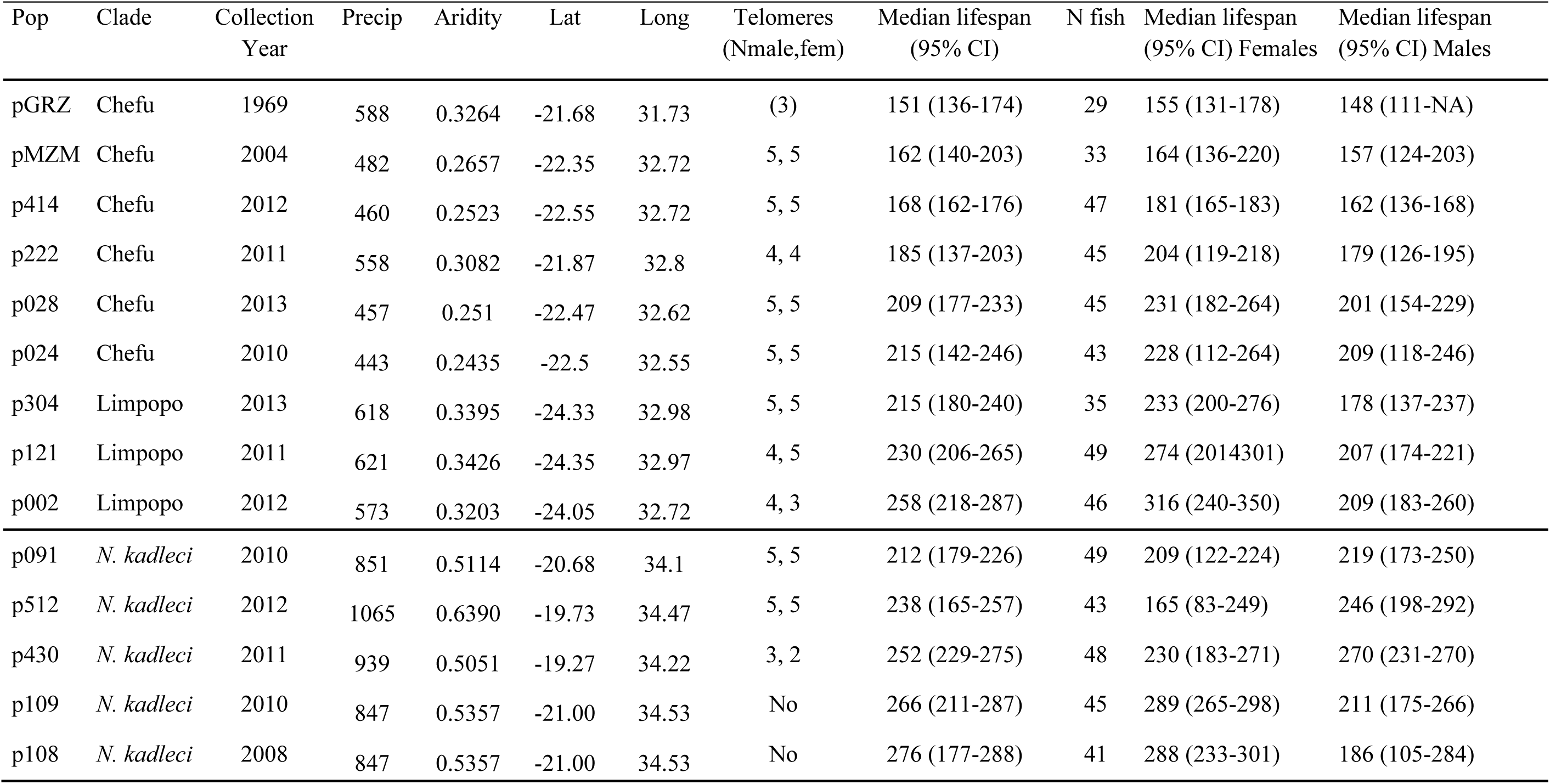
List of all strains from the *N. furzeri-N. kadleci* species complex used in the study. Populations are ordered according to median lifespan. Precipitation represents mean annual precipitation total, aridity is the index of aridity calculated from precipitation-evaporation differentials (both derived from CGIAR Consortium for Spatial Information: www.cgiar-csi.org). Sample sizes for telomere length study (telomeres) and longevity estimates (N fish) are shown. Median lifespan was calculated using the *surv* function in the *survival* package.

The *N. furzeri-kadleci* complex is composed of locally constrained populations living in small ephemeral pools which are genetically distinct from adjacent populations (Bartáková, Reichard et al. 2013). All strains used in our analyses were derived from natural populations, for which we knew the exact location of their original pool. We characterized the environmental conditions in which these natural populations lived using long-term data on precipitation (www.worldclim.org). The wild populations are distributed along a clear cline, where increasing altitude and decreasing latitude are associated with lower precipitation (Figure 1). Our comprehensive set of wild-derived strains confirmed previous conclusions using contrasts between pairs of strains (Tozzini, Dorn et al., 2013, Blažek, Polačik et al., 2017) that median lifespan correlates positively with local precipitation (*t*_12_ = 2.64, P = 0.021) for data on both sexes pooled). Interestingly, for sex-specific lifespan estimates (which were not tested in previous studies), this relationship holds in males (LM: *t*_12_ = 3.19, p = 0.008) but not females (*t*_13_ = 0.32, p = 0.752) (Figure 2). Our results indicate that males, but not females, from arid regions have shorter median lifespans than those from humid regions, suggesting that perhaps strains from arid regions would exhibit shorter telomeres.

**Fig. 1.**
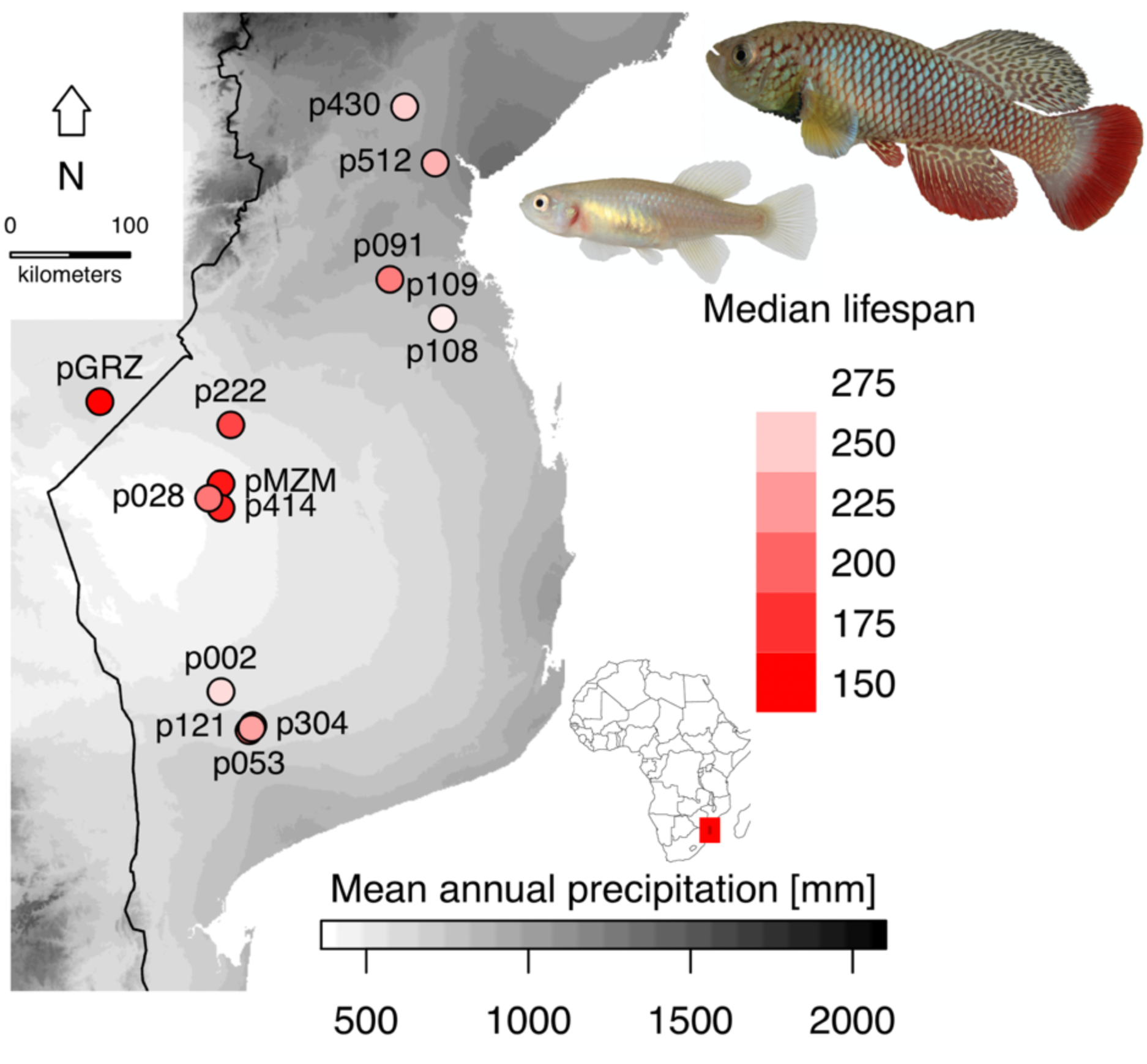
Geographic origin of study strains along the cline of precipitation, with male (larger) and female (smaller) *Nothobranchius furzeri* inset. Median lifespan (sexes pooled) is visualized by intensity of coloration for each original strain location (red: shortest lifespan; white: longest lifespan). Sex differences in fish body size are proportional.

**Fig. 2.**
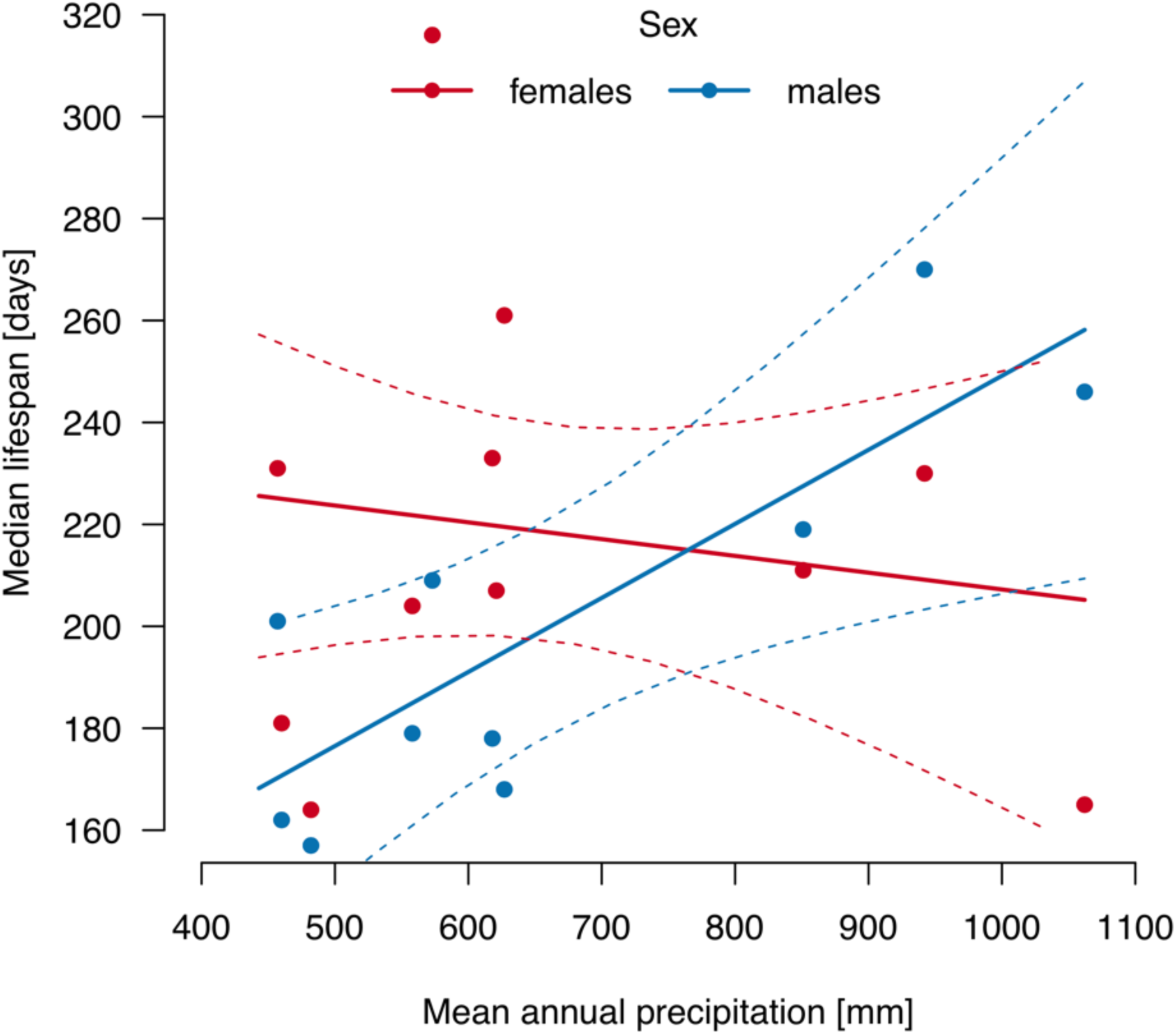
Relationship between median lifespan of males (blue) and females (red) and mean annual precipitation in their region of origin. Lines were fitted using linear regression.

### Average telomere length is a good descriptor of telomere length distribution

Telomere length was estimated using Telomere Restriction Fragment (TRF) by Southern blot analysis on samples from 11 wild derived populations (10 fish per population, with equal sex ratio). The final sample size (n = 92, Table 1) was slightly reduced due to lower quality of genomic DNA. All fish examined were young adults of the same age (28 days old), except for a single population (p222; 69 days old). All tissue samples comprised small biopsies from the caudal fin.

We first explored the relationships between various descriptors of telomere length distributions. Using a biplot based on a Principal Component Analysis (PCA) of the data distribution parameters (Figure 3), we found that disparate measures of telomere length (mean, median, 5-95% quantiles) were all highly positively correlated (Supplementary Table 1). The measures of telomere length variability within individual fish (inter-quartile range and coefficient of variation) were not strongly related to the estimates of telomere length (Figure 3, Supplementary Table 1). Based on this information, we used the mean as the measure of telomere length and the coefficient of variation (CV) as a measure of intra-individual telomere length variability in further analyses. These two measures were not correlated (Pearson, r = −0.01, n = 92, p = 0.940).

**Fig. 3.**
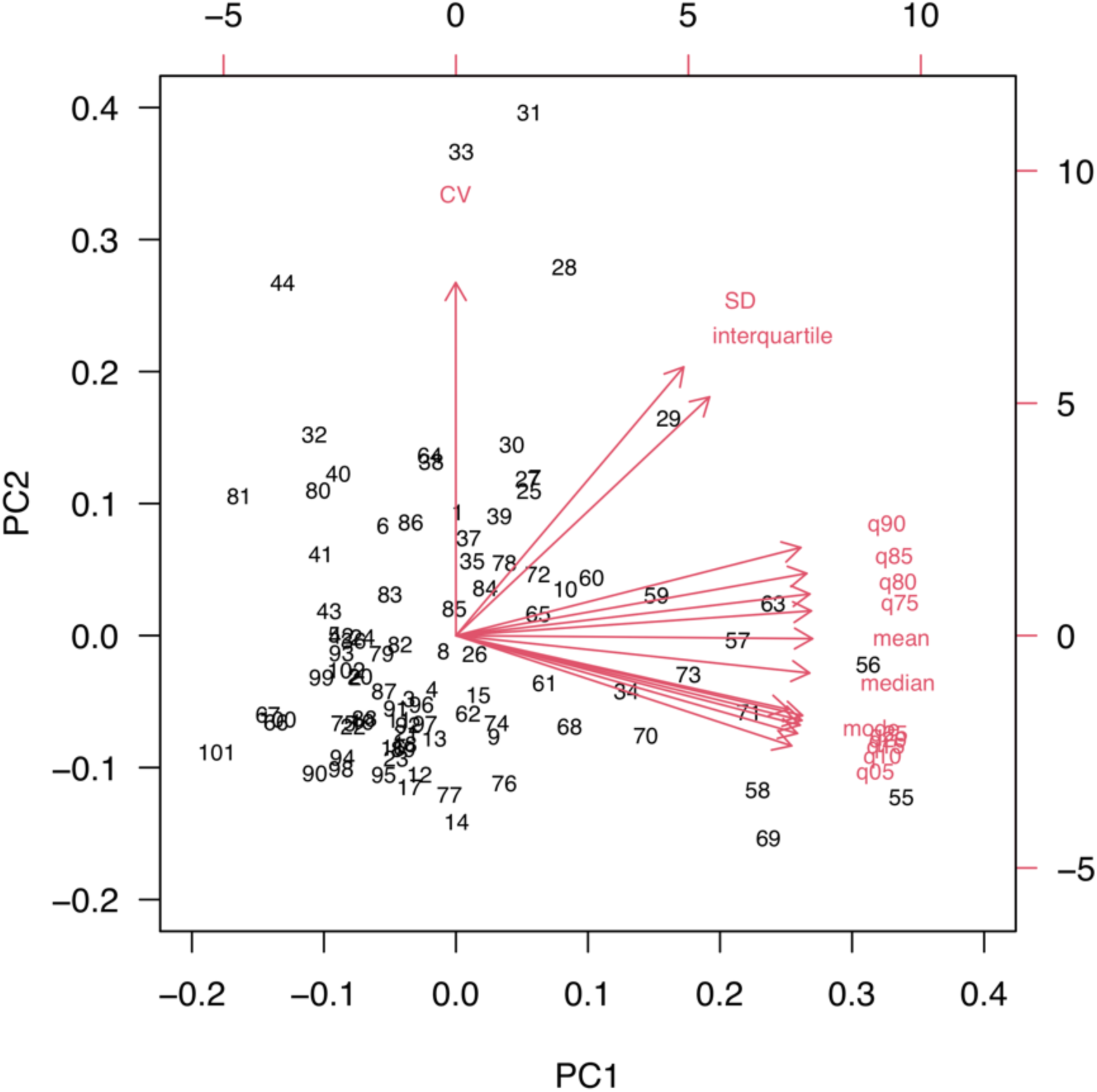
Biplot visualization of telomere length distribution parameters, based on Principal Component analysis. Vectors represent estimated parameters; the numbers denote sample ID.

### Telomere length is shorter in males and in faster growing individuals

The exploratory analysis revealed a difference in mean telomere length, from 5.29 (s.e. 0.42) to 9.92 (s.e. 0.42) kbp across various strains (Figure 4, Supplementary Figure 1, Supplementary Table 2). When comparing telomere length between males and females, we accounted for inter-strain variation using strain ID as a random effect.

**Table 2.**
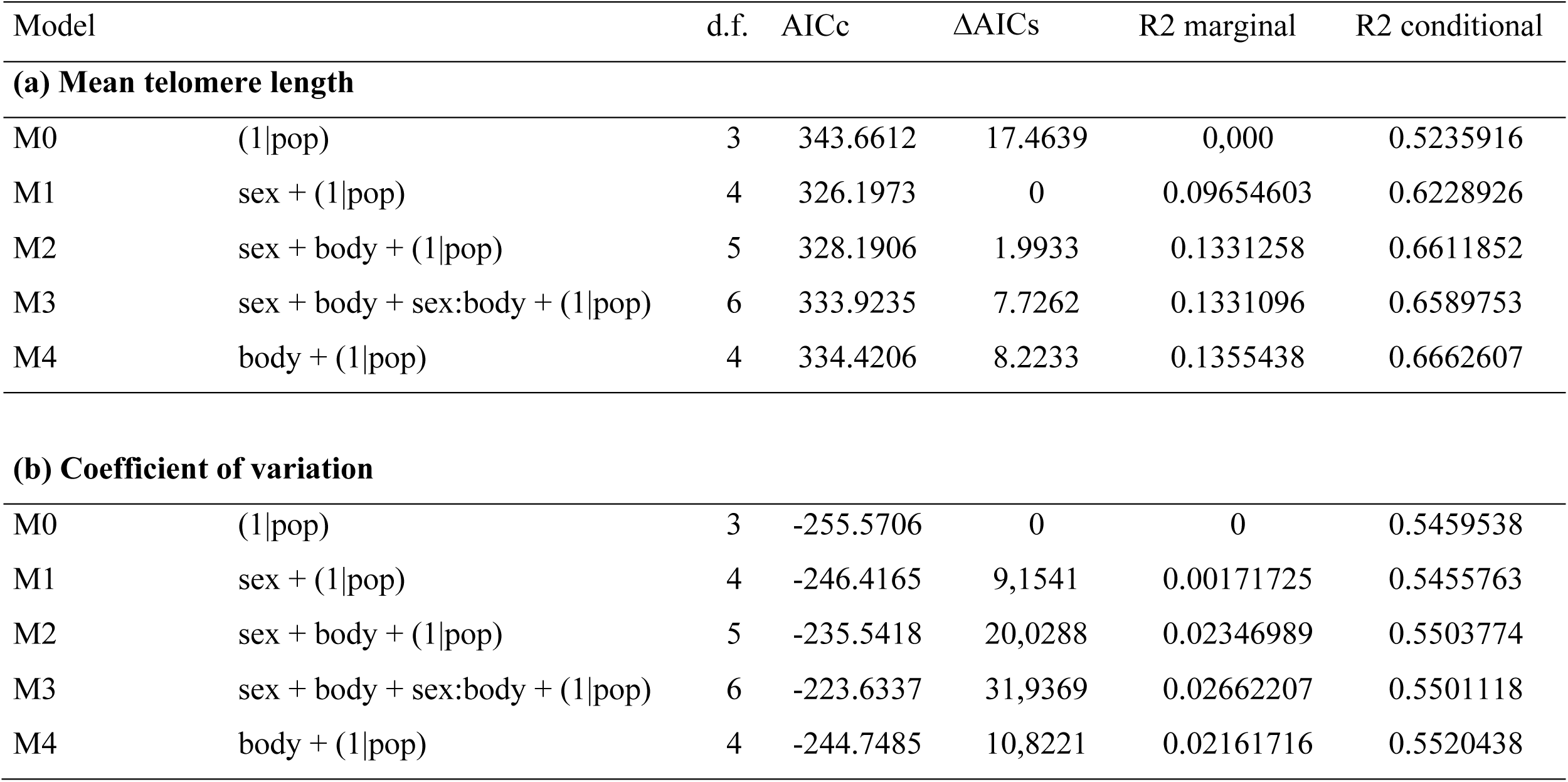
List of candidate models to test the relationship of mean telomere length (a) and its intra-individual variation (b) with sex and juvenile growth. Model structure (fixed effects and their interaction, and random effect of strain ID), degrees of freedom (d.f.), Akaike Information Criterion corrected for small sample size (AICc), difference between the best fitting model and candidate model (ΔAICc) indicating strength of support for a particular model, amount of variability explained by fixed effects (R^2^_marginal_) and by combination of fixed and random effects (R^2^_conditional_).

**Fig. 4.**
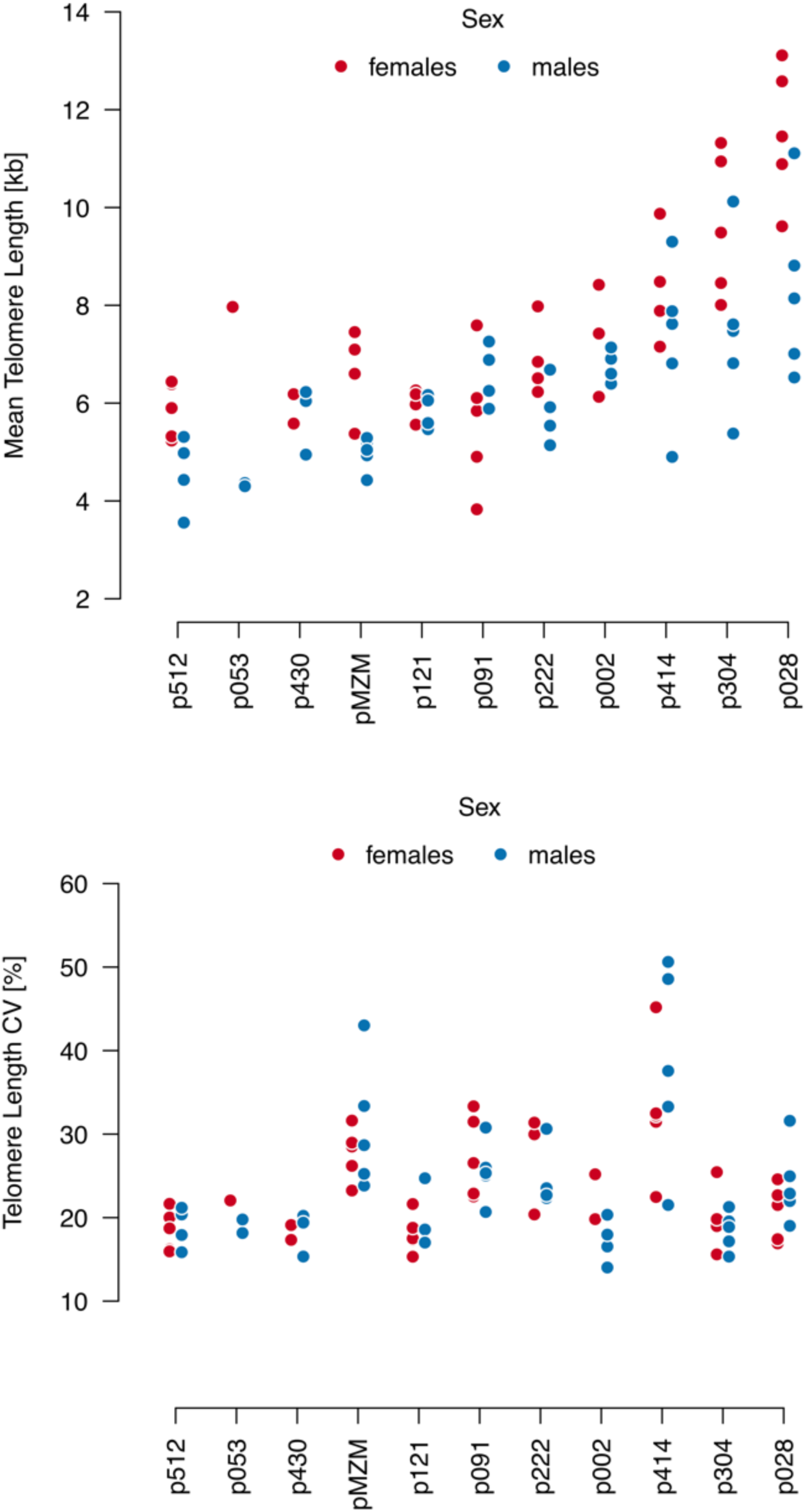
Among strains variability in (A) mean telomere length and (B) intra-individual variation in telomere length, estimated as Coefficient of Variation (CV). Strains are ranked from shortest to longest telomere length.

We first tested how parameters of telomere length vary in relation to sex and juvenile growth (measured as individual body size at age of 28 days). Our set of candidate models contained the two explanatory variables and their interaction, along with a null model (Table 2). Candidate models were fitted to the data and compared using the Akaike Information Criterion corrected for small sample size (AICc). The best model contained only sex (Table 2a) and demonstrated that males had shorter telomeres than females (Linear Mixed Model with strain ID modelled as a random factor: *F*_1,80.4_ = 23.13, P < 0.001). Across strains, mean telomere length was 7.37 kbp (s.e. 0.45) in females and 6.18 (s.e. 0.24) in males. Strain identity explained a large part of variation in all models; the values of R^2^_GLMM conditional_ were approximately 6 times larger than R^2^_GLMM marginal_ (Table 2).

We note that the second best model (model M2 in Table 2a, ΔAIC = 1.99) indicated that rapid juvenile growth (i.e. large size at the age of 28 days) was associated with relatively shorter telomere length (*F*_1,88.9_ = 4.47, P = 0.037) but the trend became weaker and statistically non-significant after removal of the strain with older (and larger) fish (p222: 69 old vs. 28 days old fish) (*F*_1,75.7_ = 3.52, P = 0.064; Supplementary Table 3).

**Table 3.** List of candidate models to test the relationship of mean telomere length (a) and its intra-individual variation (b) with precipitation. Model structure (fixed effects and their interaction, and random effect of strain ID), degrees of freedom (d.f.), Akaike Information Criterion corrected for small sample size (AICc), difference between the best fitting model and candidate model (ΔAICc) indicating strength of support for a particular model, amount of variability explained by fixed effects (R^2^_marginal_) and by combination of fixed and random effects (R^2^_conditional_).

| Model | | d.f. | AICc | $\Delta\text{AICc}$ | R2 marginal | R2 conditional |
| --- | --- | --- | --- | --- | --- | --- |
| **(a) Mean telomere length** |  |  |  |  |  |  |
| M0 | (1\|pop) | 3 | 343.6612 | -8.341 | 0 | 0.5235916 |
| M1 | precip + (1\|pop) | 4 | 352.6750 | -17.3548 | 0.1741896 | 0.5407045 |
| M2 | precip + sex + (1\|pop) | 5 | 335.3202 | 0 | 0.2649685 | 0.6376086 |
| M3 | precip + sex + precip:sex + (1\|pop) | 6 | 344.3068 | -8.9866 | 0.2844145 | 0.6560993 |
| **(b) Coefficient of variation** |  |  |  |  |  |  |
| M0 | (1\|pop) | 3 | -255.5706 | 0 | 0 | 0.5459538 |
| M1 | precip + (1\|pop) | 4 | -238.9254 | 16.6452 | 0.1275769 | 0.5670824 |
| M2 | precip + sex + (1\|pop) | 5 | -229.7255 | 25.8451 | 0.1286001 | 0.5665069 |
| M3 | precip + sex + precip:sex + (1\|pop) | 6 | -211.1557 | 44.4149 | 0.1348708 | 0.5715629 |

Intra-individual variation in telomere length was not related to sex, juvenile growth or their interaction (Table 2b).

### Telomere length tends to be shorter in strains from more humid environments

We then tested whether total annual precipitation in the original habitat, which is associated with (male) lifespan, also correlated with telomere length. The best fitting model (Table 3a) included sex differences in telomere length (*F*_1,80.5_ = 23.08, P < 0.001) and a trend towards negative association with precipitation totals (Figure 5; *F*_1,9.0_ = 4.11, P = 0.072), suggesting that killifish strains from regions with higher precipitation tend to possess shorter telomeres.

**Fig. 5.**
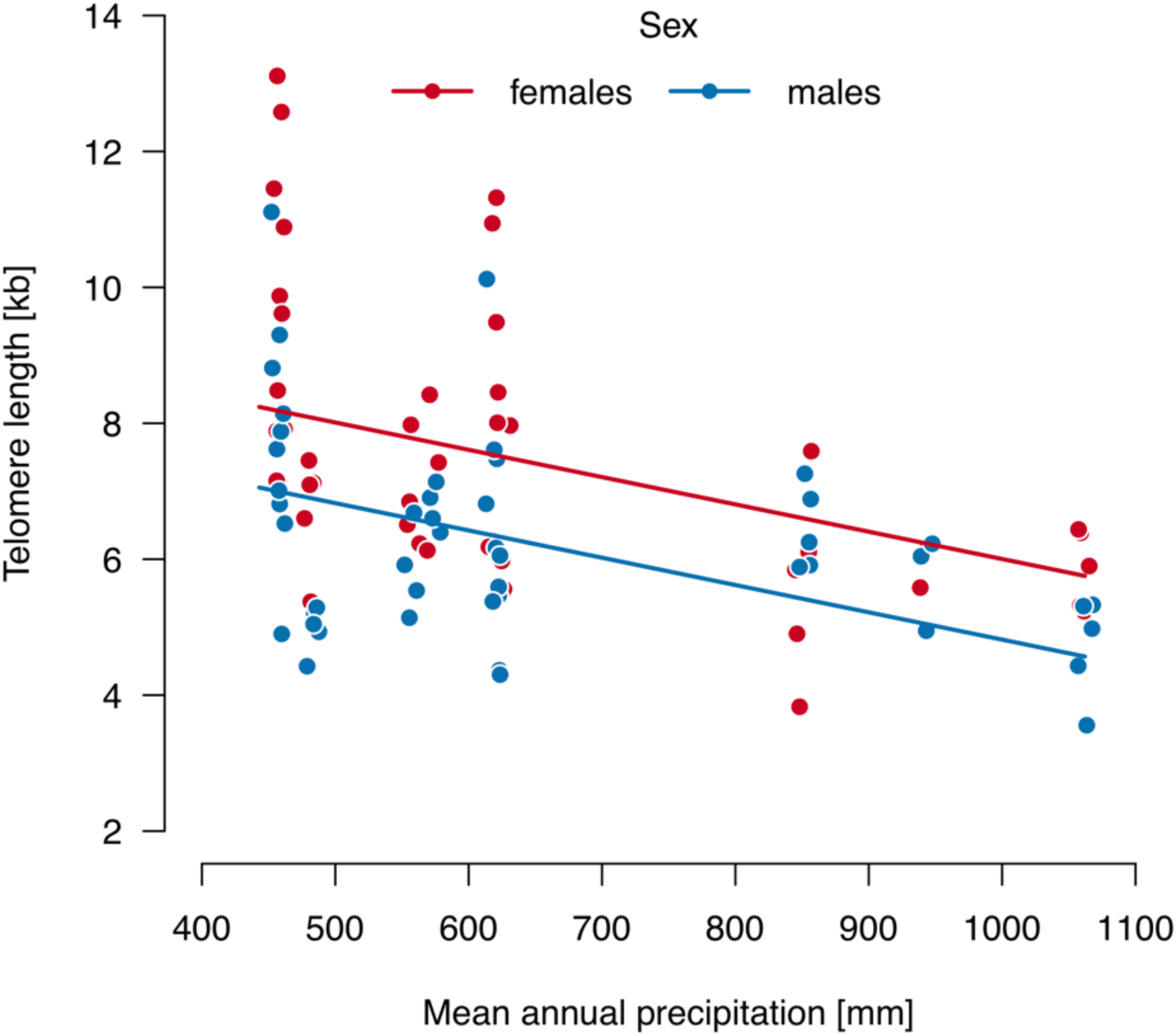
Relationship between mean telomere length and mean annual precipitation in male (blue) and female (red) killifish. Solid lines represent relationship fitted with a mixed model including sex and precipitation as fixed factors and strain ID as a random factor.

Telomere length variability (CV) was neither related to precipitation nor to sex (Table 3b), although it varied considerably across strains (Figure 4b).

### Telomere length is not associated with median lifespan

Given that the sexes differed in their lifespans, we used sex-specific lifespan estimates to test the association between telomere length and population-level longevity (Figure 6). The best model (Table 4a) confirmed sex differences in telomere length (*F*_1,8071_ = 18.57, P < 0.001), but did not demonstrate a link between mean telomere length and median lifespan estimate (*F*_1,88.7_ = 2.39, P = 0.126). Therefore, consistent with previous studies at the inter-specific level, telomere length does not appear to predict longevity among different killifish strains.

**Table 4.**
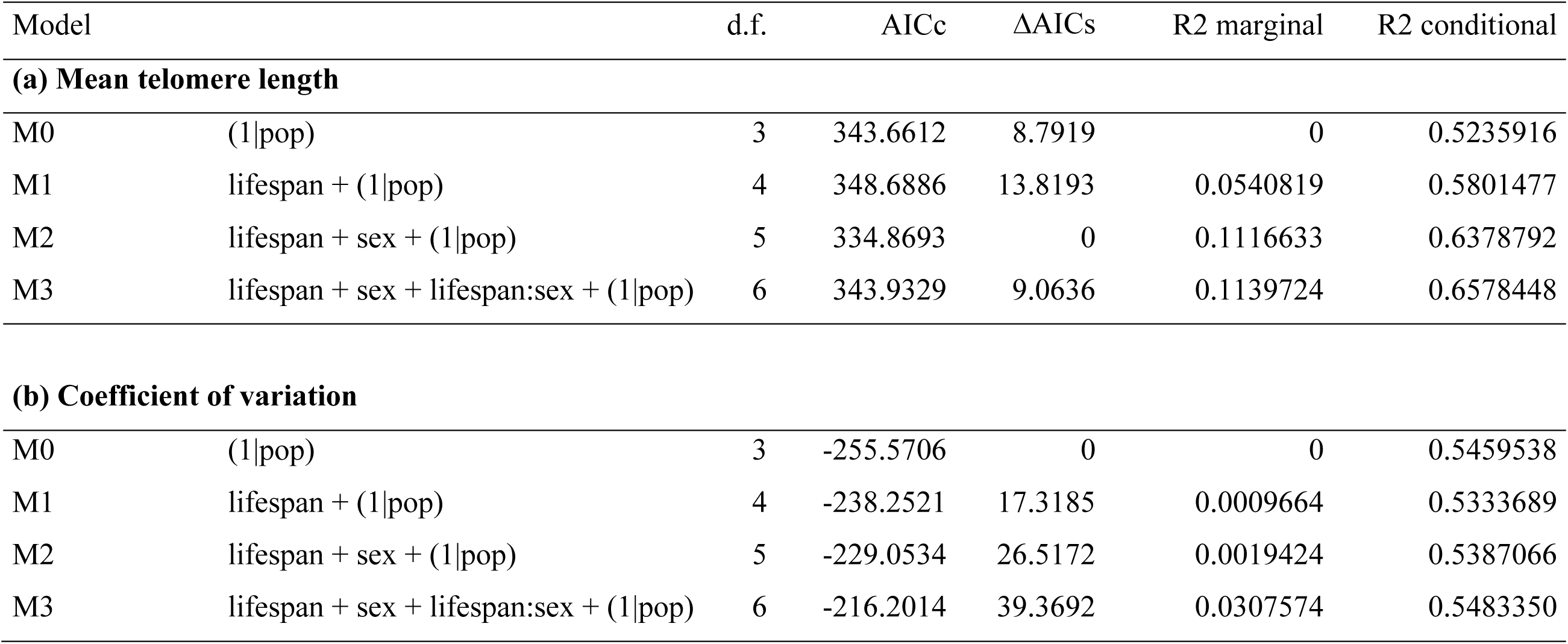
List of candidate models to test the relationship of mean telomere length (a) and its intra-individual variation (b) with median sex-specific strain-level lifespan estimates. Model structure (fixed effects and their interaction, and random effect of strain ID), degrees of freedom (d.f.), Akaike Information Criterion corrected for small sample size (AICc), difference between the best fitting model and candidate model (ΔAICc) indicating strength of support for a particular model, amount of variability explained by fixed effects (R^2^_marginal_) and by combination of fixed and random effects (R^2^_conditional_).

| Model | | d.f. | AICc | $\Delta\text{AICs}$ | R2 marginal | R2 conditional |
| --- | --- | --- | --- | --- | --- | --- |
| **(a) Mean telomere length** |  |  |  |  |  |  |
| M0 | (1\|pop) | 3 | 343.6612 | 8.7919 | 0 | 0.5235916 |
| M1 | lifespan + (1\|pop) | 4 | 348.6886 | 13.8193 | 0.0540819 | 0.5801477 |
| M2 | lifespan + sex + (1\|pop) | 5 | 334.8693 | 0 | 0.1116633 | 0.6378792 |
| M3 | lifespan + sex + lifespan:sex + (1\|pop) | 6 | 343.9329 | 9.0636 | 0.1139724 | 0.6578448 |
| **(b) Coefficient of variation** |  |  |  |  |  |  |
| M0 | (1\|pop) | 3 | -255.5706 | 0 | 0 | 0.5459538 |
| M1 | lifespan + (1\|pop) | 4 | -238.2521 | 17.3185 | 0.0009664 | 0.5333689 |
| M2 | lifespan + sex + (1\|pop) | 5 | -229.0534 | 26.5172 | 0.0019424 | 0.5387066 |
| M3 | lifespan + sex + lifespan:sex + (1\|pop) | 6 | -216.2014 | 39.3692 | 0.0307574 | 0.5483350 |

**Fig. 6.**
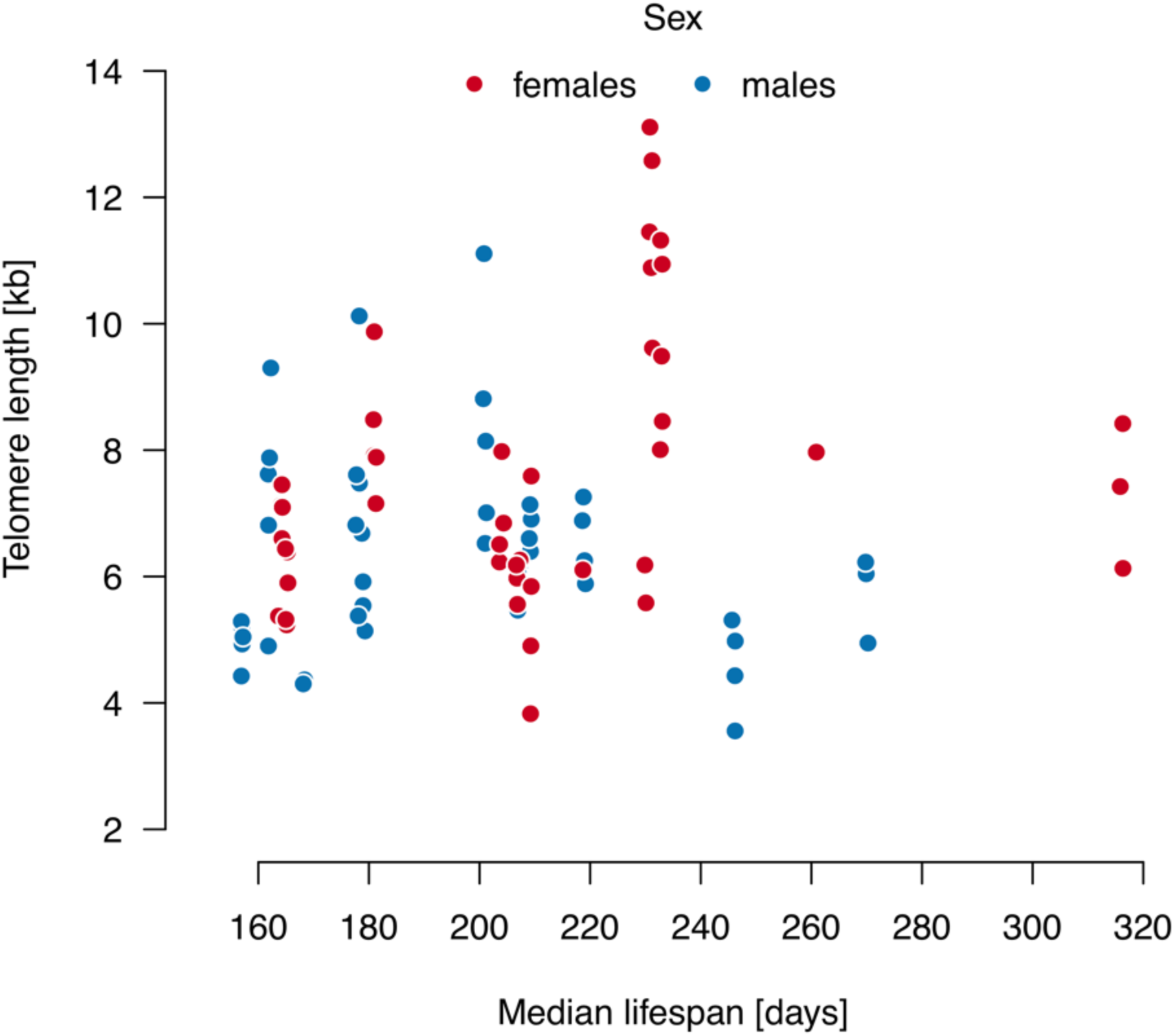
Relationship between telomere length and sex-specific strain-level lifespan estimates.

For telomere length variability (CV), none of the candidate models was better than a null model (Table 4b).

### Telomere length in other *Nothobranchius* species

Finally, we expanded telomere length analysis to three other *Nothobranchius* species from Mozambique (*N. orthonotus, N. pienaari, N. rachovii*) that widely coexist with *N. furzeri-kadleci* species complex (Reichard, Janáč et al. 2017). We also included two species from Tanzania (*N. rubripinnis, N. eggersi*), where a more humid climate is linked to a longer duration of wet habitat phase (Reichard 2015).

Telomere length among these species was comparable (range 3.6-7.3 kbp, Supplementary Figure 2) but generally lower than the *N. furzeri-N. kadleci* group. Estimated telomere length was generally longer for the relatively shorter-lived *N. orthonotus* (strain MZCS 002 Limpopo: mean (s.e.) = 5.63 (0.29), n=4; MZCS 528 Beira: 6.03 (0.35), n=2) compared to *N. pienaari* (MZCS 505 Limpopo: 4.29 (0.25), n=4; MZCS 514 Beira: 5.86 (0.29), n=3) and *N. rachovii* (MZCS 096 Beira: 3.65, n=1). Tanzanian *Nothobranchius* species had telomere lengths comparable to Mozambican species (*N. rubripinnis* T33: 5.60 (0.25), n=4; *N. eggersi* T52: 5.45 (0.35), n=2). Finally, we also tested telomere length in a highly inbred GRZ strain of *N. furzeri* that was bred under different laboratory conditions from the other fish and, therefore, was not included in the overall analysis. Mean telomere length in the GRZ fish (7.33 (0.29), n=3) corresponded to the predicted values based on their origin in a relatively dry region.

## DISCUSSION

Telomeres and telomerase prevent the continuous erosion of chromosome-ends caused by lifelong cell division. In several species, including humans, telomeres shorten with age and their decline is considered a hallmark of ageing (López-Otín, Blasco et al. 2013). Telomere structure and function is conserved across eukaryotes and their exhaustion leads to multiple effects both at the cellular and systemic level, causing disruption of organ homeostasis and disease (Armanios, Alder et al. 2009, Carneiro, Henriques et al. 2016). Traditionally, telomere length has been considered indicative of “biological age” and a useful proxy to predict individual health status and longevity. However, evidence from several non-human vertebrates now contradicts this assumption as simplistic.

In our large sample of wild-derived annual killifish strains obtained from a gradient of environmental conditions and natural lifespans, we found that telomere length varied across strains and tended to be shorter in populations that came from more humid regions where populations naturally live longer. The natural habitats of annual killifish are seasonal pools in the African savannah. They are inundated during the rainy season, fish hatch, grow rapidly and attain sexual maturity within a few weeks (Vrtílek, Žák et al. 2018a). The pools desiccate during the dry season and killifish populations persist as dormant eggs in the pool sediment. The wet phase may vary from a few weeks to several months (Vrtílek, Žák et al. 2018b) depending on the amount of precipitation in the area (Blažek, Polačik et al. 2017). Habitat duration, which varies considerably across the precipitation gradient of killifish distribution, determines maximum lifespans. Male lifespans in our laboratory assays were strongly associated with annual precipitation levels in their area of their origin, though female lifespans were not. Consistent with previous observations across species (Gomes, Ryder et al. 2011), killifish from more humid regions (and hence with longer lifespans, both predicted and observed) had shorter telomeres than those from arid regions. This complements an analogous trend found across mammalian species (Gomes, Ryder et al. 2011) and demonstrates that telomere length differences among populations reflect patterns found among species.

Alternatively, telomere length differences across strains may simply have been fixed through genetic drift. Natural *Nothobranchius* populations are genetically distinct units (Bartáková, Reichard et al. 2015) but are more fragmented and have smaller effective population sizes in the dry part of their range (Willemsen, Cui et al. 2020). Killifish populations from dry regions experience increased mutation load, leading to a reduction in their lifespans (Cui, Medeiros et al. 2019, Willemsen, Cui et al. 2020). However, our data demonstrate that dry-region populations tend to have longer telomeres, including the highly inbred GRZ population with apparently the strongest mutation load. This trend is further strengthened by the telomere length of other three *Nothobranchius* species from Mozambique (*N. orthonotus, N. pienaari, N. rachovii*), widely coexisting with the *N. furzeri-kadleci* species complex and two species from a more humid region in coastal Tanzania (*N. eggersi, N. rubripinnis*). Telomere length was comparable in all these longer-lived species (Blažek, Polačik et al. 2017, Wildekamp 2004) but generally shorter than in the *N. furzeri-N. kadleci* complex, suggesting that genetic drift is unlikely a driver of such consistent population-specific differences in telomere length.

Sex differences in telomere length were consistent across strains (Supplementary Table 2), with male telomeres 22% shorter on average. Remarkably, males also had consistently shorter lifespans than females in all nine *N. furzeri* (but only in 2 of 5 *N. kadleci*) populations. All our experimental fish were housed in social tanks. This pattern of sex differences in lifespan corroborates conclusions from a separate study (Reichard, Blažek et al. 2020) where *N. furzeri* (but not *N. kadleci*) males in social tanks suffered higher baseline mortality than females. Higher male mortality was not reflected in sex differences in any measure of functional aging (oxidative stress, kidney and liver histopathology, accumulation of lipofuscin) (Reichard, Blažek et al. 2020). Individual variation in telomere length distribution did not differ between the sexes, discounting the possibility that females possess mechanisms for countering cells with short telomeres. However, it is plausible that males are more susceptible to telomere erosion due to higher hormonal and environmental stress. *Nothobranchius* fish display remarkable sexual dimorphism and dichromatism. Males are larger and more brightly colored than females and continuously compete for mating opportunities (Cellerino, Valenzano et al., 2016, Reichard and Polačik 2019). Male reproductive strategy (higher growth, investment in bright coloration and aggressive interactions) therefore implicates a higher metabolic cost and more rapid growth than the female reproductive strategy (Bonduriansky, Maklakov et al. 2008). This may underlie the faster decline in telomere length in males, which we recorded as a shorter telomere length in young adult males.

Even within the sexes, rapid juvenile growth tended to be associated with relatively shorter telomere length. Our sampling at the age of 28 days encompassed the period of most rapid growth rate in *Nothobranchius* lifespan, with fish growing from an initial body length of 5 mm to 50 mm (Blažek, Polačik et al. 2013, Vrtílek, Žák et al. 2018a). This raises a plausible hypothesis that growth rate, primarily in early life, at least partly affects the kinetics of telomere erosion and overall telomere length. We now plan a follow-up study with repeated longitudinal sampling of the same individuals to test this hypothesis.

Studies on telomere length in the field of ecology and evolution typically use real-time PCR assays to determine Telomere-to-Single Copy Gene (T/S) ratio which provides an indirect estimate of mean telomere length (Lai, Wright et al. 2018). However, the detrimental effects of telomere shortening are expected to arise from a frequently dividing subpopulation of cells within a particular tissue, which may not be well represented in the T/S ratio. We estimated telomere length using the robust TRF method that yields a direct distribution of telomere fragments in the sample. Importantly, we confirmed that estimates of mean values are a useful proxy for telomere length distribution as they closely correlated with other parameters of telomere length distribution, including a set of 5-95% quantiles. Thus, mean telomere length represents a suitable estimate of telomere kinetics in adult *Nothobranchius* killifishes. Additionally, the measure of telomere length variability within an individual, the coefficient of variation, was not associated with any pattern in our data (between sexes, among strains, or in juvenile growth). This suggests that the dispersion in telomere length distribution is a strain-specific character controlled by regulatory pathways other than those used in determining telomere length. As telomeres shorten, wider telomere distributions will give rise to more critically short telomeres than narrow ones, thus providing an independent parameter to control telomere decline with age.

Overall, our results demonstrate a clear variation in telomere length between sexes and environments among wild-derived strains of *Nothobranchius* fishes and highlight their suitability for understanding the adaptive value of telomere length across individuals, populations, and species.

## MATERIALS AND METHODS

### Fish origin and environmental data

Study strains were descendants of wild populations imported from Mozambique and the adjacent area of Zimbabwe. In this region, a clear gradient of aridity is generated by a decrease in total precipitation with increasing distance from the coast, increasing altitude and decreasing latitude associated with lower precipitation (Figure 1; Blažek, Polačik et al. 2017). Fish from wild populations were imported between 2009 and 2012 during dedicated expeditions and wild-derived strains were maintained in the Institute of Vertebrate Biology (IVB) in Brno, Czech Republic according to published protocol (Polačik, Blažek et al. 2016). One strain (GRZ) was imported in 1969 from Zimbabwe and another strain (MZM-0410, coded as pMZM in our dataset) was imported in 2004. The origin of each study strain is illustrated in Figure 1 and GPS coordinates are provided in Table 1. Other strains of Mozambican (*N. orthonotus, N. pienaari, N. rachovii*) and Tanzanian species (*N. eggersi and N. rubripinnis*) originate from our own imports between 2008 and 2017 (Supplementary Table 4). Housing conditions were the same for all species.

Data on precipitation (represented by annual rainfall totals) and aridity were downloaded from www.worldclim.org. Aridity was measured as aridity index (total precipitation/model-estimated evapotranspiration). Given that precipitation and aridity were highly collinear (Pearson r = 0.989, n = 12, P < 0.001), we only present analyses using total precipitation.

### Fish housing and lifespan estimates

Fish were kept at the IVB facility under standard conditions and care was taken to minimize inbreeding (i.e. a large number of fish were sources of offspring for the next generation, though no specific breeding design was followed).

For the estimates of lifespan, eggs of 14 strains were hatched simultaneously. The eggs were kept in diapause at 19°C. Two weeks prior to hatching, the eggs were placed in 28°C to initiate release from Diapause II and development to prehatching stage. In August 2016, the incubation substrate (peat) was watered with dechlorinated tap water (conductivity 600microS/cm, temperature 16°C) in a 2L tank. Fish hatched within 1 day and were immediately fed on freshly hatched *Artemia* nauplii. After 5 days, fish were moved to larger tanks (40 L) and their density was standardized across strains. At 3 weeks, fish were weaned from *Artemia* nauplii to frozen chironomid larvae, a standard diet of captive *Nothobranchius* (Polačik, Blažek et al. 2016).

At 6 weeks, fish density was standardized to 50 fish per strain, though some strains had a slightly lower density (Table 1). All tanks were monitored daily and any dead fish was recorded and removed. Daily censuses continued until all fish were dead.

Median lifespan was calculated for each strain (and each sex within a strain) using the *survfit* function in the *survival* package. 95% confidence intervals for median lifespan estimates were calculated using a log-log function.

### Telomere length estimates

Telomere length was estimated through Telomere Restriction Fragment (TRF) analysis followed by Southern blotting. Fish from 12 strains (all strains used in lifespan estimate except for GRZ, p108, and p109) were hatched on 6 May 2019, following the same protocol as for lifespan estimates. When all fish were sexually mature, at the age of 28 days, a biopsy from the caudal fin was taken from 5 male and 5 female fish from each population and flash frozen in liquid nitrogen. Samples of other species were handled in the same way, except they were stored in 100% ethanol instead of being flash frozen. Due to logistic reasons, one population (*N. furzeri*, MZCS 222) was hatched earlier (26 March 2019) and was older (69 days) at the time of sampling, although it was raised following identical methods and kept in the same room. To account for the potential effect of age on telomere length, we re-ran statistical analyses with this population excluded from the dataset. This population had no effect on the qualitative interpretation and, in the main text, we present the final analysis with this population included. The exception is the test of relationship between body length at sexual maturity (28 days) and telomere length, where we present both the full and truncated datasets as this is where the removal of MZCS 222 strain affected the strength of statistical association.

Genomic DNA was extracted using a lysis buffer (Thermo Scientific #K0512) supplemented with 1 mg/ml Proteinase K (0.5 mg/ml final concentration, Sigma, MO, USA) and RNase A (1:100 dilution, Sigma, MO, USA). Samples were incubated at 50°C for 18 h in a thermomixer. Purification of genomic DNA was carried out using equilibrated phenol-chloroform (Sigma, MO, USA) and chloroform-isoamyl alcohol extraction (Sigma, MO, USA). Following quantification and normalization, genomic DNA was digested with RsaI and HinfI enzymes (NEB, MA, USA) for 12 h at 37°C. Digested samples were electrophoresed on a 20 cm 0.6% agarose gel, in 0.5% TBE buffer, at 4°C for 17 h at 110 constant voltage. Southern blotting was performed as previously described (Kimura, Stone et al. 2010) with minor changes; a telomere probe, (CCCTAA)_4_, labelled with [α-32P]-dCTP through the Prime-it II random primer labelling kit (Stratagene) was used. Raw data from Southern Blots are presented as Supplementary Figure 1.

### Statistical analysis

We first extracted pre-selected parameters of the telomere distribution and analyzed them using Principal Component Analysis (PCA) to test their covariance. We then selected two parameters (mean telomere length and coefficient of variation) representing two uncorrelated descriptors of individual-fish telomere populations for further analyses.

To test the effects of predictors on telomere length and intra-individual variability, we used an information theoretic approach (Burnham and Anderson 2002). A set of biologically plausible models was constructed, with various level of complexity, and including a null (intercept-only) model. All models included strain identity as a random effect to control for non-independence of telomere length across individuals from the same population. We then compared the fit of the models using the Akaike Information Criterion corrected for small sample size (AICc). The best model minimizes AICc value. Model within ΔAICc between 2 and 5 are considered to have a comparable power.

Analyses were performed in *lmer* and *MuMIn* packages in the R 3.5.2 environment. Before applying statistical models, data exploration was undertaken as recommended by Zuur and Ieno et al. (2016).

## Supporting information

Set of Supplementary Tables and Figures

## Acknowledgement

Funding came from Czech Science Foundation (19-01781S) (to MR), Portuguese Fundação para a Ciência e Tecnologia (POCI-01-0145-FEDER-016390) (to MGF) and French Université Côte d’Azur Academie 4 Installation Grant (to MGF). All work was carried out in accordance with relevant guidelines and regulations. Collection of wild strains complied with legal regulations of Mozambique (collection licenses: DPPM/053/7.10/08, 175/154/ IIP/2009/DARPE, DPPM/083/7.10/10, DPPM/330/7.10/10, DPPM/069/7.10/11, DPPM/088/7.10/12). Experimental work was approved by the Ethical Committee of the Institute of Vertebrate Biology (No. 163-12) and by Ministry of Agriculture (CZ 62760203) in accordance with legal regulations of the Czech Republic. We thank R. Spence for valuable comments and English corrections. All primary data are available at Figshare repository (doi: 10.6084/m9.figshare.12888566).

## Author contributions

MR and MGF conceived and designed the study. RB and MP collected data on lifespan. MV and MR raised fish for telomere length estimates. KG, TF performed genomic DNA extractions, TRF and Southern hybridizations. KG, TF and MGF analyzed and interpreted telomere length. MR and MV conducted statistical analyses. MR, KG and MGF drafted the manuscript. All authors significantly contributed to the final text.

