## Supplementary material for "Environment and sex control lifespan and telomere length in wild-derived African killifish": Set of Supplementary Tables and Figures

**Supplementary Figure 1**. Representative image of TRF analysis of genomic DNA by Southern Blot (random primer-labelled telomeric probe $(CCCTAA)_4$ $^{32}P$-dCTP). Each fish is marked with a unique number from 1 to 30, numbers in red indicate females and numbers in blue indicate males. Fish from different populations are labelled by codes p222 to p512. ND= non digested control; L=Ladder (size of ladder lanes is indicated in Kb on the left); X= samples not analyzed due to either poor genomic DNA quality or incomplete digestion, not included in the analyses.

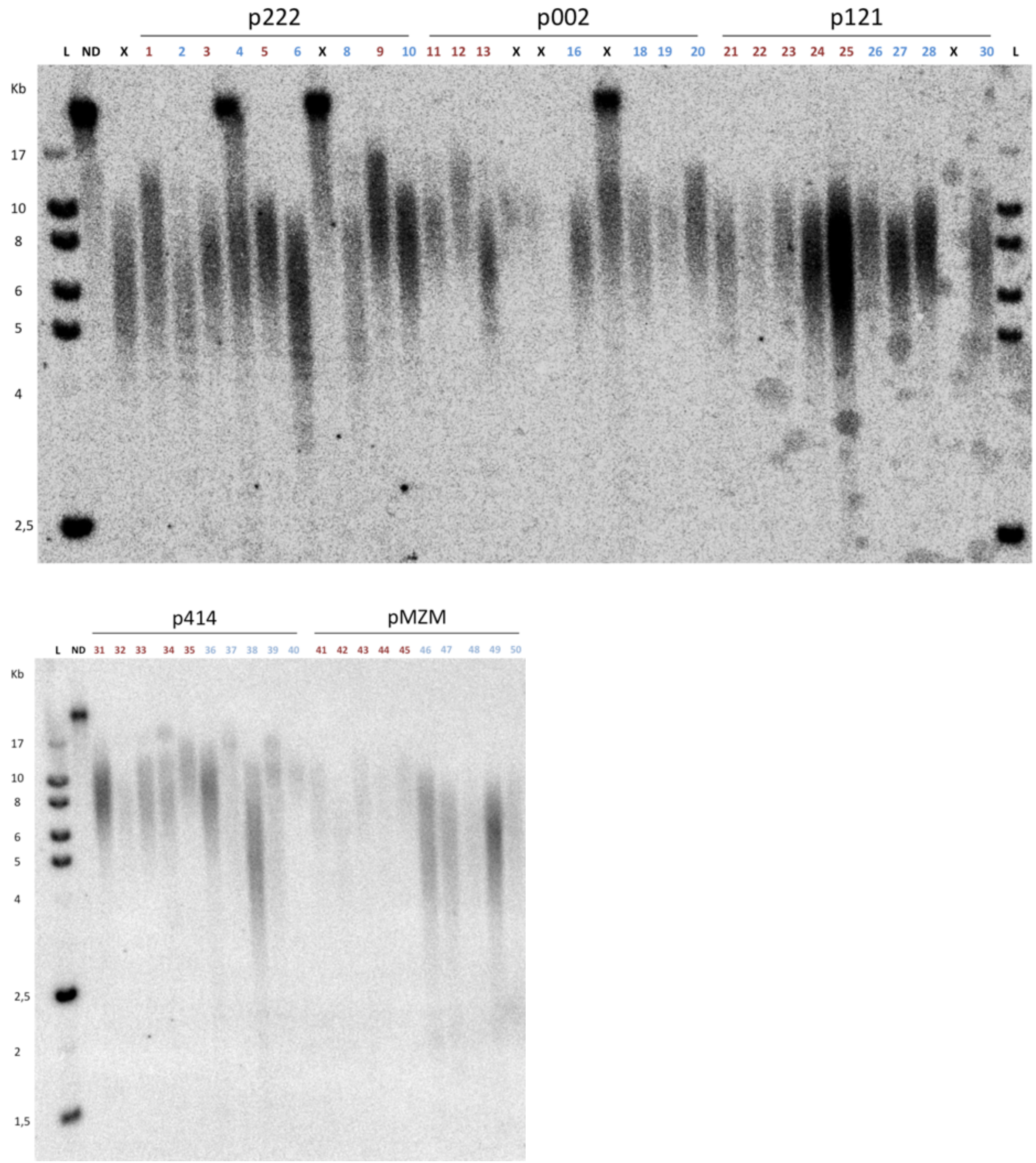


(continues on next page)

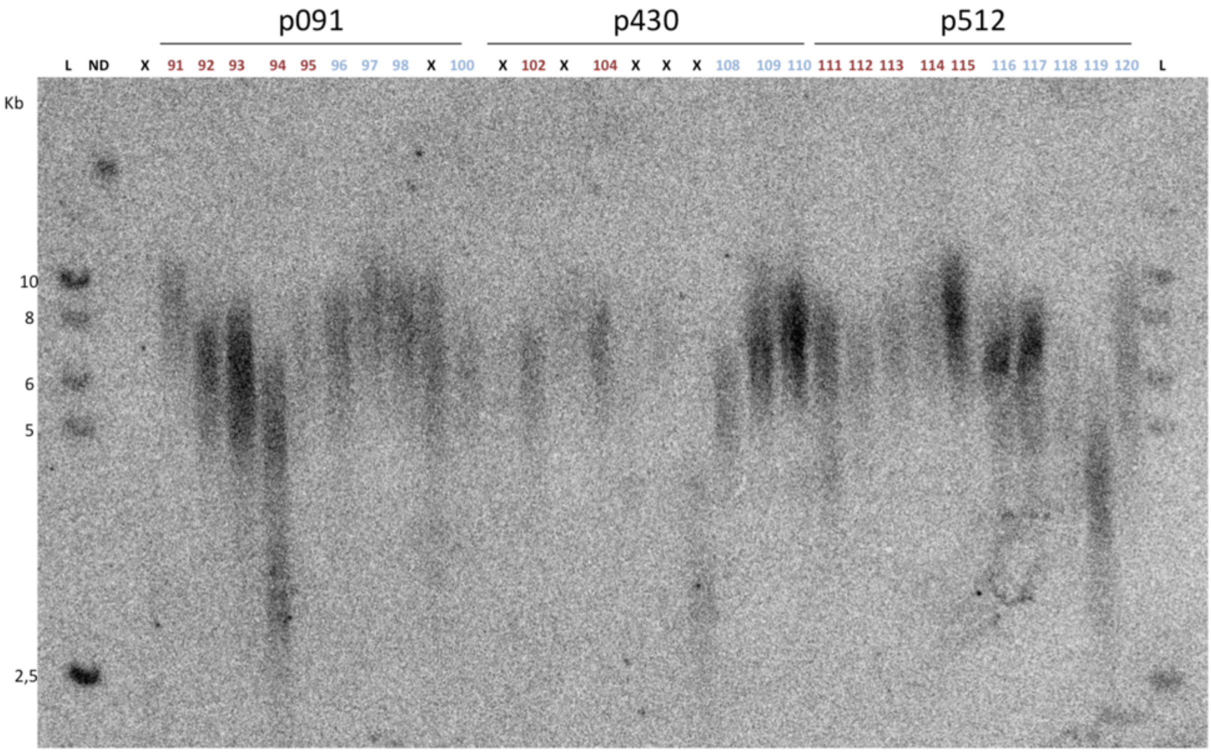
p091
p430
p512
L ND X 91 92 93 94 95 96 97 98 X 100 X 102 X 104 X X X 108 109 110 111 112 113 114 115 116 117 118 119 120 L
Kb
10
8
6
5
2,5

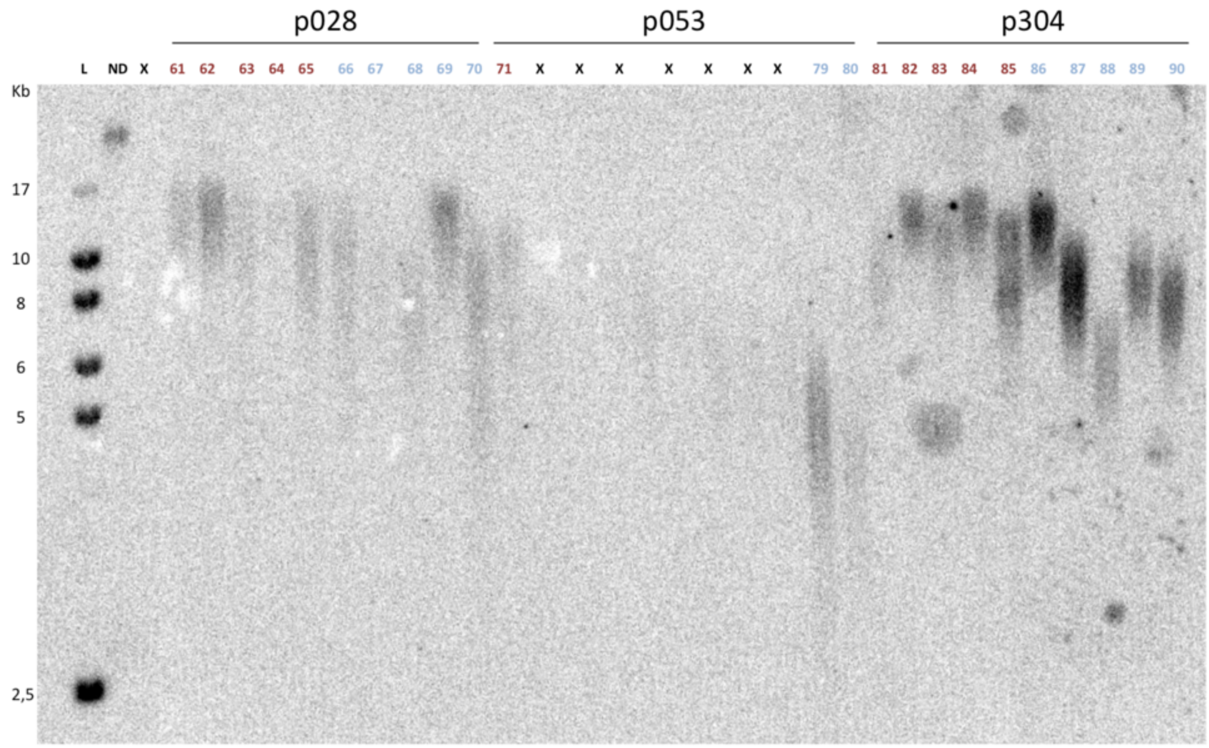
p028
p053
p304
L ND X 61 62 63 64 65 66 67 68 69 70 71 X X X X X X X 79 80 81 82 83 84 85 86 87 88 89 90
Kb
17
10
8
6
5
2,5

**Supplementary Figure 2.** Variation in mean telomere length among strains from *Nothobranchius* species other than *N. furzeri* and *N. kadleci* and in GRZ strain of *N. furzeri* bred under different conditions. Each symbol represents one individual. For codes and species, see Supplementary Table 6.

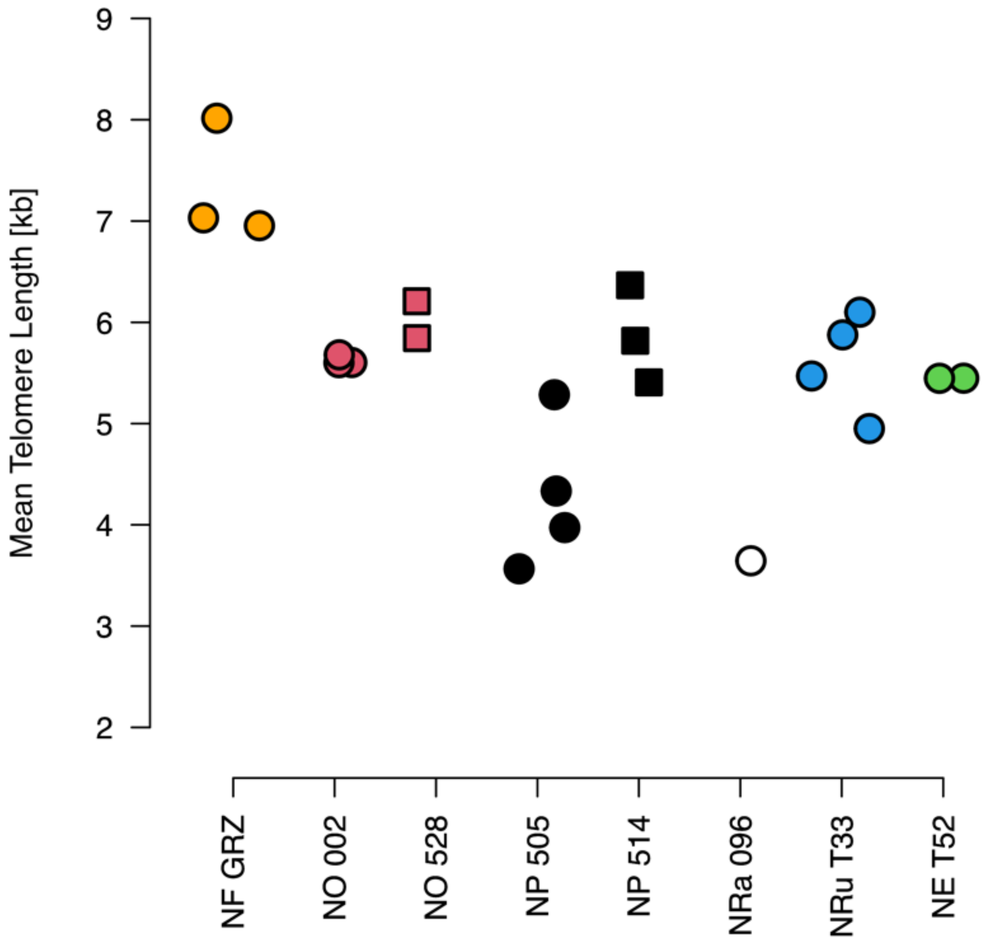

**Supplementary Figure 3.** Kaplan-Meier visualization of survival in study strains, plotted separately for males (blue) and females (red).

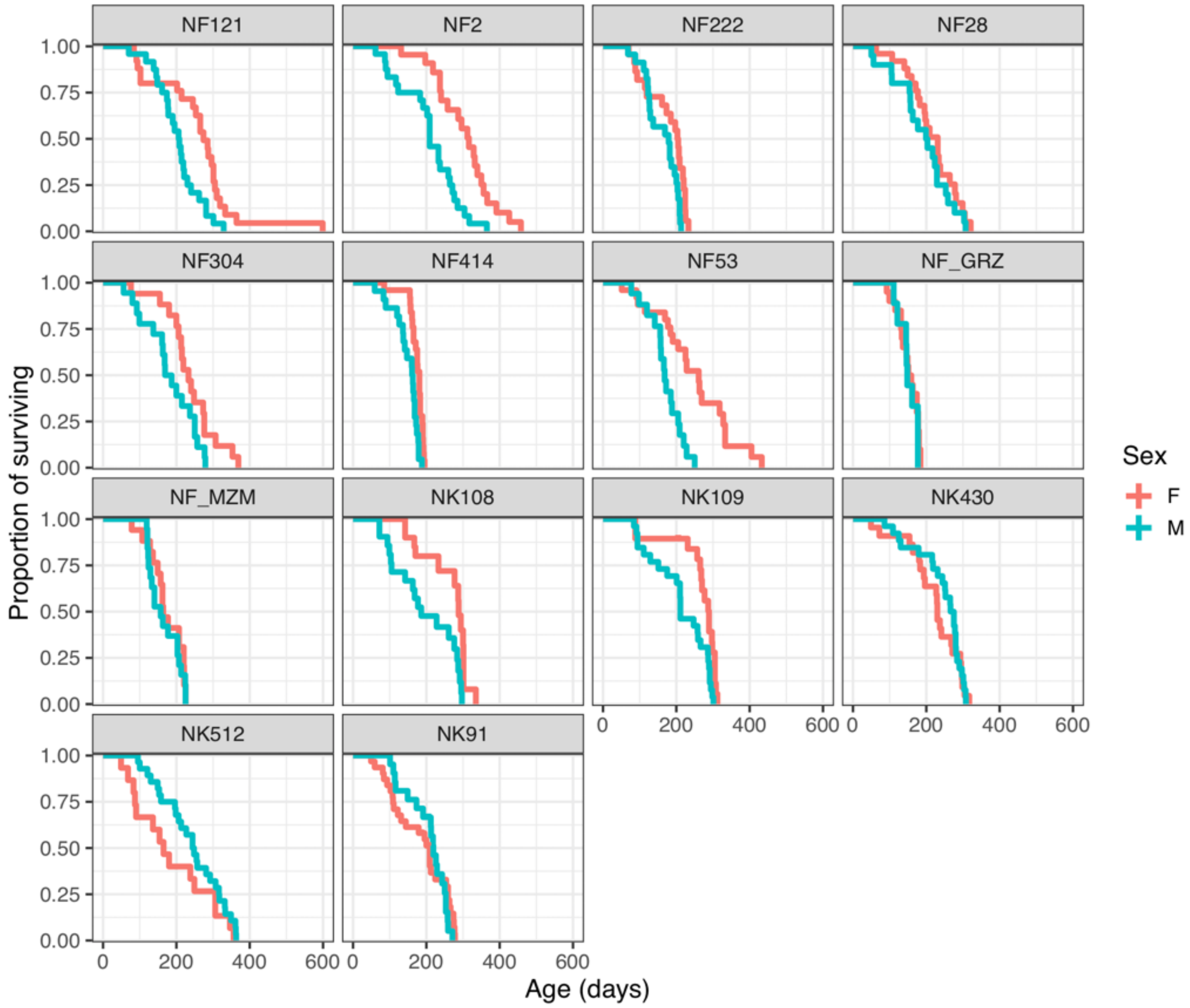

**Supplementary Table 1.** Correlation matrix of telomere distribution data parameters, with estimates of centrality (mean, median, mode), quantiles (5, 15, 25, 75, 85, 90%) and variability (interquartile range: IQ_range; Standard Deviation: SD; Coefficient of Variation: CV). For visualization, see Supplementary Figure 2.

| | Mean | Median | Mode | q05 | q15 | q25 | q75 | q85 | q90 | IQ range | SD |
|---|---|---|---|---|---|---|---|---|---|---|---|
| Median | 0.993 | | | | | | | | | | |
| Mode | 0.929 | 0.950 | | | | | | | | | |
| q05 | 0.944 | 0.961 | 0.913 | | | | | | | | |
| q15 | 0.966 | 0.979 | 0.938 | 0.991 | | | | | | | |
| q25 | 0.975 | 0.989 | 0.948 | 0.983 | 0.993 | | | | | | |
| q75 | 0.995 | 0.983 | 0.913 | 0.914 | 0.941 | 0.952 | | | | | |
| q85 | 0.982 | 0.956 | 0.872 | 0.872 | 0.905 | 0.917 | 0.993 | | | | |
| q90 | 0.964 | 0.931 | 0.841 | 0.834 | 0.871 | 0.885 | 0.979 | 0.996 | | | |
| IQ range | 0.701 | 0.635 | 0.528 | 0.458 | 0.513 | 0.529 | 0.762 | 0.818 | 0.849 | | |
| SD | 0.633 | 0.550 | 0.440 | 0.369 | 0.430 | 0.455 | 0.682 | 0.757 | 0.807 | 0.931 | |
| CV | -0.008 | -0.103 | -0.206 | -0.300 | -0.245 | -0.217 | 0.065 | 0.168 | 0.240 | 0.642 | 0.747 |

**Supplementary Table 2.** Mean (standard error) estimates of telomere length for each strain for data on both sexes pooled, and separately for males and females. Strains are ordered according to median lifespan.

| Population | Pooled | | Males | | Females | |
|---|---|---|---|---|---|---|
| pMZM | 5.85 | (0.424) | 4.98 | (0.469) | 6.73 | (0.522) |
| p414 | 7.78 | (0.424) | 7.30 | (0.469) | 8.26 | (0.522) |
| p222 | 6.36 | (0.474) | 5.82 | (0.525) | 6.89 | (0.583) |
| p053 | 5.54 | (0.774) | 4.33 | (0.742) | 7.97 | (1.166) |
| p028 | 9.92 | (0.424) | 8.32 | (0.469) | 11.53 | (0.522) |
| p304 | 8.56 | (0.424) | 7.48 | (0.469) | 9.64 | (0.522) |
| p121 | 5.87 | (0.447) | 5.82 | (0.525) | 5.91 | (0.522) |
| p002 | 7.00 | (0.506) | 6.76 | (0.525) | 7.32 | (0.673) |
| p091 | 6.05 | (0.424) | 6.44 | (0.469) | 5.65 | (0.522) |
| p512 | 5.29 | (0.424) | 4.72 | (0.469) | 5.86 | (0.522) |
| p430 | 5.80 | (0.599) | 5.74 | (0.606) | 5.88 | (0.825) |

**Supplementary Table 3.** List of candidate models to test the relationship of mean telomere length (a) and its intra-individual variation (b) with sex and juvenile growth for the dataset with strain 222 (containing older fish) excluded. Model structure (fixed effects and their interaction, and random effect of population identity), degrees of freedom (d.f.), Akaike Information Criterion corrected for small sample size (AICc), difference between the best fitting model and candidate model (ΔAICc) indicating strength of support for a particular model, amount of variability explained by fixed effects ($R^2_{marginal}$) and by combination of fixed and random effects ($R^2_{conditional}$).

| Model | | d.f. | AICc | ΔAICs | R2 marginal | R2 conditional |
|---|---|---|---|---|---|---|
| **(a) Mean telomere length** | | | | | | |
| M0 | (1\|pop) | 3 | 318.7904 | 15.0907 | 0 | 0.5356554 |
| M1 | sex + (1\|pop) | 4 | 303.6997 | 0 | 0.09114431 | 0.628192 |
| M2 | sex + body + (1\|pop) | 5 | 306.3782 | 2.6785 | 0.1087655 | 0.6539556 |
| M3 | sex + body + sex:body + (1\|pop) | 6 | 310.9822 | 7.2825 | 0.1078984 | 0.6512705 |
| M4 | body + (1\|pop) | 4 | 311.7669 | 8.0672 | 0.08056567 | 0.6341793 |
| **(b) Coefficient of variation** | | | | | | |
| M0 | (1\|pop) | 3 | -231.0669 | 0 | 0 | 0.5678328 |
| M1 | sex + (1\|pop) | 4 | -222.1130 | 8,9539 | 0.002414862 | 0.5681472 |
| M2 | sex + body + (1\|pop) | 5 | -212.5577 | 18,5092 | 0.0229539 | 0.5746474 |
| M3 | sex + body + sex:body + (1\|pop) | 6 | -223.6337 | 7,4332 | 0.02662207 | 0.5501118 |
| M4 | body + (1\|pop) | 4 | -221.6326 | 9,4343 | 0.02164824 | 0.5781961 |

**Supplementary Table 6.** List of *Nothobranchius* strains from clades other than the *N. furzeri-N. kadleci* species complex. Precipitation represents mean annual precipitation totals. Aridity is the index of aridity calculated from precipitation-evaporation differentials. Latitude and longitude for the original location of each strain is given. Sample size for the telomere length study (N) is also indicated.

| Species | Population | Collection | Precipitation | Aridity | Lat | Long | N |
|---|---|---|---|---|---|---|---|
| *N. orthonotus* | p2 | 2012 | 573 | 0.320 | -24.05 | 32.72 | 4 |
| *N. orthonotus* | p528 | 2012 | 1269 | 0.812 | -19.68 | 34.77 | 2 |
| *N. pienaari* | p505 | 2012 | 490 | 0.274 | -23.52 | 32.57 | 4 |
| *N. pienaari* | p514 | 2012 | 1275 | 0.818 | -19.68 | 34.78 | 3 |
| *N. rachovii* | p96 | 2008 | 1385 | 0.930 | -19.80 | 34.90 | 1 |
| *N. eggersi* | t52 | 2017 | 1353 | 0.861 | 6.482 | 38.914 | 2 |
| *N. rubripinnis* | t33 | 2017 | 924 | 0.650 | 7.211 | 39.175 | 5 |
